# Random Tactile Noise Stimulation Reveals Beta-Rhythmic Impulse Response Function of the Somatosensory System

**DOI:** 10.1101/2022.09.03.506453

**Authors:** Samson Chota, Rufin VanRullen, Rasa Gulbinaite

## Abstract

Both passive tactile stimulation and motor actions result in dynamic changes in beta-band (15-30 Hz Hz) oscillations over somatosensory cortex. Similar to alpha-band (8-12 Hz) power decrease in the visual system, beta-band power also decreases following stimulation of the somatosensory system. This relative suppression of alpha and beta oscillations is generally interpreted as an increase in cortical excitability. Here, next to traditional single-pulse stimuli, we employed a random intensity continuous right index finger tactile stimulation (white noise), which enabled us to uncover an impulse response function (IRF) of the somatosensory system. Contrary to previous findings, we demonstrate a burst-like initial increase rather than decrease of beta activity following white noise stimulation (human participants, N = 18, 8 female). These beta bursts, on average, lasted for 3 cycles and their frequency was correlated with resonant frequency of somatosensory cortex, as measured by a multi-frequency steady-state somatosensory evoked potential (SSSEP) paradigm. Furthermore, beta-band bursts shared spectro-temporal characteristics with evoked and resting-state beta oscillations. Taken together, our findings not only reveal a novel oscillatory signature of somatosensory processing that mimics the previously reported visual IRFs, but also point to a common oscillatory generator underlying spontaneous beta bursts in the absence of tactile stimulation and phase-locked beta bursts following stimulation, the frequency of which is determined by the resonance properties of the somatosensory system.

**Significance Statement:** The investigation of the transient nature of oscillations has gained great popularity in recent years. The findings of bursting activity rather than sustained oscillations in the beta-band has provided important insights into its role in movement planning, working memory, inhibition and reactivation of neural ensembles. In this study, we show that also in response to tactile stimulation the somatosensory system responds with ∼3 cycle oscillatory beta-band bursts, whose spectro-temporal characteristics are shared with evoked and resting-state beta-band oscillatory signatures of the somatosensory system. As similar bursts have been observed in the visual domain, these oscillatory signatures might reflect an important supramodal mechanism in sensory processing.

## Introduction

Oscillatory dynamics are assumed to play a critical role in the processing of perceptual information (Buzsaki, 2006), with different sensory modalities operating preferentially in different frequency bands (Kayser, 2019; Klimesch et al., 2007; Spitzer & Haegens, 2017). In response to tactile stimulation, as well as during movement preparation and execution, the power of beta oscillations (15-30 Hz) over somatosensory cortices decreases (Hari et al., 1997; Neuper & Pfurtscheller, 2001; Parkkonen et al., 2015; Salenius et al., 1997; Salmelin & Hari, 1994). This reduction in beta-band power has generally been interpreted as an increase in excitability, allowing the somatosensory system to process external and interoceptive information during perception, motor coordination and feedback (Cassim et al., 2000, 2001; Gaetz & Cheyne, 2006). Hence, beta oscillations might have a similar role to that of alpha oscillations in the visual system, namely cortical inhibition (Bonnefond & Jensen, 2012; Engel & Fries, 2010; Haegens et al., 2011; Händel et al., 2011; Jensen & Mazaheri, 2010; Jones et al., 2010; Klimesch et al., 2007; Linkenkaer-Hansen et al., 2004; Shin et al., 2017).

The interpretation of alpha and beta as inhibitory brain rhythms is based on observations of *sustained* (∼500 ms duration) power decreases in trial-average waveforms following the stimulus (Hari et al., 1997; Neuper & Pfurtscheller, 2001; Parkkonen et al., 2015). A more nuanced spatiotemporal dynamic and functional role of low-frequency rhythms has been revealed by focusing on their transient nature. Single-trial analyses, in case of beta oscillations in the somatosensory system, revealed a short-lived (∼150 ms, 3 cycles) burst-like behavior of beta oscillations (Little et al., 2019; Lundqvist et al., 2018; Shin et al., 2017). Similar oscillatory transients have been uncovered in the visual system using reverse correlation between visual input and output (∼10 Hz, lasting up to 1 sec) (VanRullen & Macdonald, 2012). Both of these approaches revealed a transient increase in oscillatory power hidden in global sustained decreases in power, which suggests a more active than traditionally assumed and temporally-precise role of these rhythms, e.g. in sequence learning and predictive processing (Alamia & VanRullen, 2019; Huang et al., 2018).

In this study, we aimed to investigate the transient nature and frequency tuning of beta oscillations in the somatosensory system using several complementary approaches: (1) tactile random white noise stimulation (all frequencies in a 1-75 Hz range), which allowed us to compute participant-specific impulse response function (IRF) of the somatosensory system; (2) frequency-by-frequency rhythmic tactile stimulation in a 12-39 Hz frequency range (steady-state somatosensory evoked potentials, SSSEPs), which allowed us to estimate the resonance, or maximal response amplitude, frequency of somatosensory cortex for each participant; (3) pulsed tactile stimulation – a typical event-related potential protocol; and (4) spontaneous pre-stimulus beta-band activity. This broad approach allowed us to compare multiple beta oscillatory signatures within somatosensory cortex and thus provide comprehensive evidence for the link between all of them.

We found that somatosensory IRFs, just like their alpha-rhythmic visual counterparts, contained beta-frequency components and were unique to each participant in frequency (M = 24.78 Hz; SD = 6.18 Hz). Tactile stimulation elicited beta bursts that lasted on average for ∼3 cycles (M = 2.81 cycles; SD = 0.35 cycles). This is in line with reports of transient 3-cycle long movement-related beta bursts (Little et al., 2019; Lundqvist et al., 2016). The peak frequency of somatosensory IRFs was strongly correlated with the somatosensory resonance frequency determined using an SSSEP approach, as well as with the peak frequency of stimulus-evoked and spontaneous beta oscillations.

Our findings point to a common oscillatory generator underlying spontaneous beta bursts in the absence of tactile stimulation and phase-locked beta bursts following stimulation, the frequency of which is determined by the resonance properties of the somatosensory system. We propose an active role of beta-band bursts in processing tactile information.

## Materials and Methods

### 1. Participants

18 participants (aged 22-41, 8 females) with normal or corrected to normal vision enrolled in the study. The number of participants was chosen based on previous literature investigating somatosensory resonance (Moungou, 2016: N=12; Müller, 2011: N=10; Snyder, 1996: N=17; Tobimatsu, 1999: N=10). Informed consent forms were signed before the experiment. The experiment was carried out in accordance with the protocol approved by the Centre National de la Recherche Scientifique (CNRS) ethical committee and followed the Code of Ethics of the World Medical Association (Declaration of Helsinki). Subjects were compensated with 10 Euro/hour.

### 2. Experimental Design

Tactile stimulation was delivered via vibrotactile electromagnetic solenoid-type actuator (Dancer Design, UK) driven by an audio amplifier connected to a microcontroller board (Figure 1A). We used amplitude-modulated stimulation using 150 Hz carrier frequency. Such approach is commonly used to study human touch perception, as time-varying stimulation strongly drives several types of mechanoreceptors (Moungou et al., 2016).

**Figure 1.**
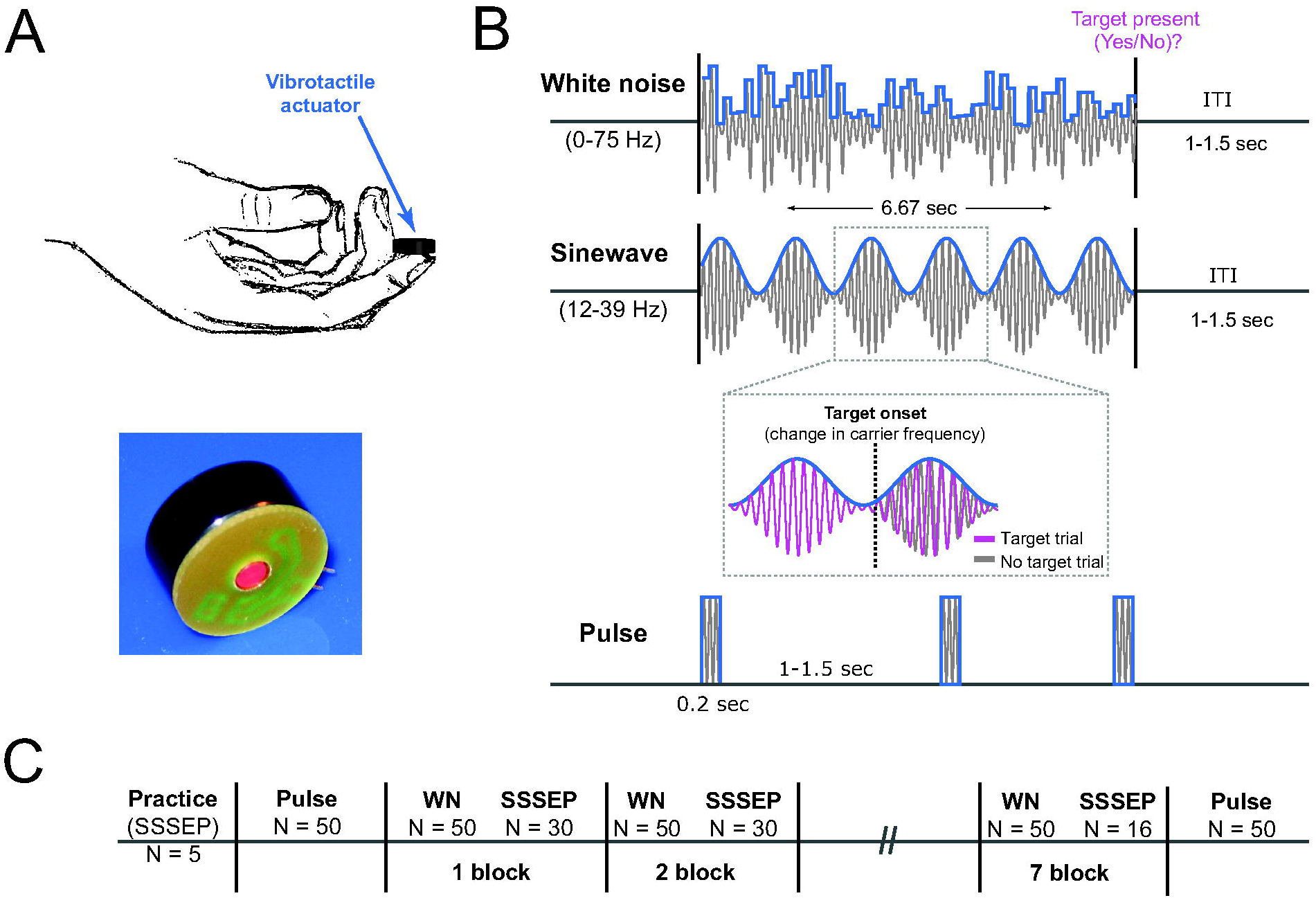
Stimulation protocol and task. **A**. Tactile stimulation was applied to participants’ right index finger with a vibrotactile actuator. **B**. Three protocols were used: 1. White noise (WN) stimulation was created by random amplitude modulation of the 150 Hz carrier sinewave. The spectral content of white noise sequences was flattened between 0 and 75 Hz. 2. SSSEP stimulation was created by rhythmic amplitude modulation of the 150 Hz carrier sinewave at one specific frequency on each trial (12 – 39 Hz). Perceptual targets were embedded in both WN and SSSEP sequences by decreasing the frequency of the carrier sinewave from 150 Hz to 127.5 Hz. **3**. Single-pulse stimulation consisted of 200 ms fixed amplitude increases of a carrier 150 Hz wave.

We used three vibrotactile stimulation protocols: (1) random white noise stimulation, (2) steady-state rhythmic stimulation, and (3) short pulses (see Figure 1B). Tactile stimulation was applied to the participants’ right index finger. White noise sequences were created by randomly modulating the amplitude of a 150 Hz carrier sinewave every cycle (i.e., every 1000/150 = 6.7 ms). The amplitude modulated signal was created by generating a random number series and normalizing the amplitude of its Fourier components before applying an inverse Fourier transform. Steady state stimuli were created by amplitude-modulating a 150 Hz carrier frequency with a constant frequency sine-wave signal at one of the frequencies. In total, we used 18 stimulation frequencies covering the beta-band frequency range and a few frequencies below and above: 12 to 20 Hz in steps of 2 Hz, 21 to 30 Hz in steps of 1 Hz, and 33 to 39 Hz in steps of 3 Hz. Each frequency was presented on 12 trials. The order of frequencies was randomized across trials.

Prior to the experiment, participants performed a practice session to become acquainted with a sensation of vibrotactile stimulation and target detection task. The task involved detecting a 1 second long 15% decrease in carrier frequency (from 150 Hz to 127.5 Hz). At the end of each trial, participants had to press with their left hand the appropriate keyboard button indicating whether the trial contained a target or not. The responses were not speeded and only 20% of trials contained targets.

Trial length for both white noise and sine-wave stimulation was 6.7 seconds (6.6625 s). The practice session consisted of 5 sine-wave stimulation trials all containing targets. After the practice session, participants were first presented with 200 pulse stimuli (200 ms duration, 150 Hz vibration) separated by a random ITI of 1-1.5 sec. Thereafter participants completed 616 experimental trials: 400 trials of white noise sequences and 216 trials of sine-wave stimulation. Experimental trials were divided into 8 blocks, with 80 trials each (7 blocks of 80 trials and 56 trials in the last block). The experimental part ended with an additional 200 single-pulse stimuli. To make sure that participants could not perform the target detection task by using auditory information (vibrotactile electromagnetic solenoid-type stimulators make a faint yet audible sound), they were using earplugs.

### 3. Data acquisition and preprocessing

We recorded participants’ EEG using a 64 channel ActiveTwo Biosemi system. Data analysis was performed in Matlab using the Fieldtrip toolbox and custom written Matlab scripts. Prior to all preprocessing steps, we visually identified and removed noisy channels. The EEG data was then re-referenced to the channel average, bandpass filtered between 1 and 200 Hz and line noise was removed using a DFT-filter (50, 100 and 150 Hz). We performed independent component analysis to identify and remove eye-movement related artifacts. Thereafter, the data was epoched for SSSEP (−1000 to 8000 ms relative to stimulus onset), white noise sequences (−1000 ms to 6625 ms relative to stimulus onset) and pulse stimulation (−400 to 600 ms relative to stimulus onset) trials separately. Baseline correction was applied by subtracting average values calculated from -1000 ms to 0 ms (WN), -200 to ms (SSSEP), or -400 ms to 0 ms (ERP) from individual trials. For SSSEP analysis, EEG data was high-pass filtered at 0.5 Hz and down-sampled to 512 Hz. Based on visual inspection, trials containing large muscle and head-movement related artifacts were removed.

### 4. Statistical Analysis

#### 4.1 ERP analysis

Event-related potentials were calculated by averaging epochs from 400 ms before to 600 ms after stimulus presentation and applying absolute baseline correction (baseline window -200 ms to 0 ms).

#### 4.2 Impulse Response Functions

Impulse response functions (IRF) were calculated by first cross-correlating the z-scored single-trial EEG signal with the white noise sequence presented at that particular trial and then averaging the result over trials. EEG trials were down-sampled to 150 Hz before cross-correlation to match the rate of vibrotactile stimulation amplitude changes (Figure 2). To evaluate statistical significance of IRFs we created a null hypothesis distribution by cross-correlating EEG signal with white noise sequences from random trials (surrogate impulse response function). This randomization procedure was repeated 10.000 times and the resulting trial-averaged IRFs served as the distribution of expected correlation values under the null-hypothesis (H0: EEG and white noise are uncorrelated). All stimulus time points were entered in the cross-correlation except the first 500 ms and the last 1000 ms to avoid influences of the onset and offset ERP to the estimate of tactile IRF. The cross-correlation was calculated for lags between -300 ms and 500 ms as follows:

**Figure 2.**
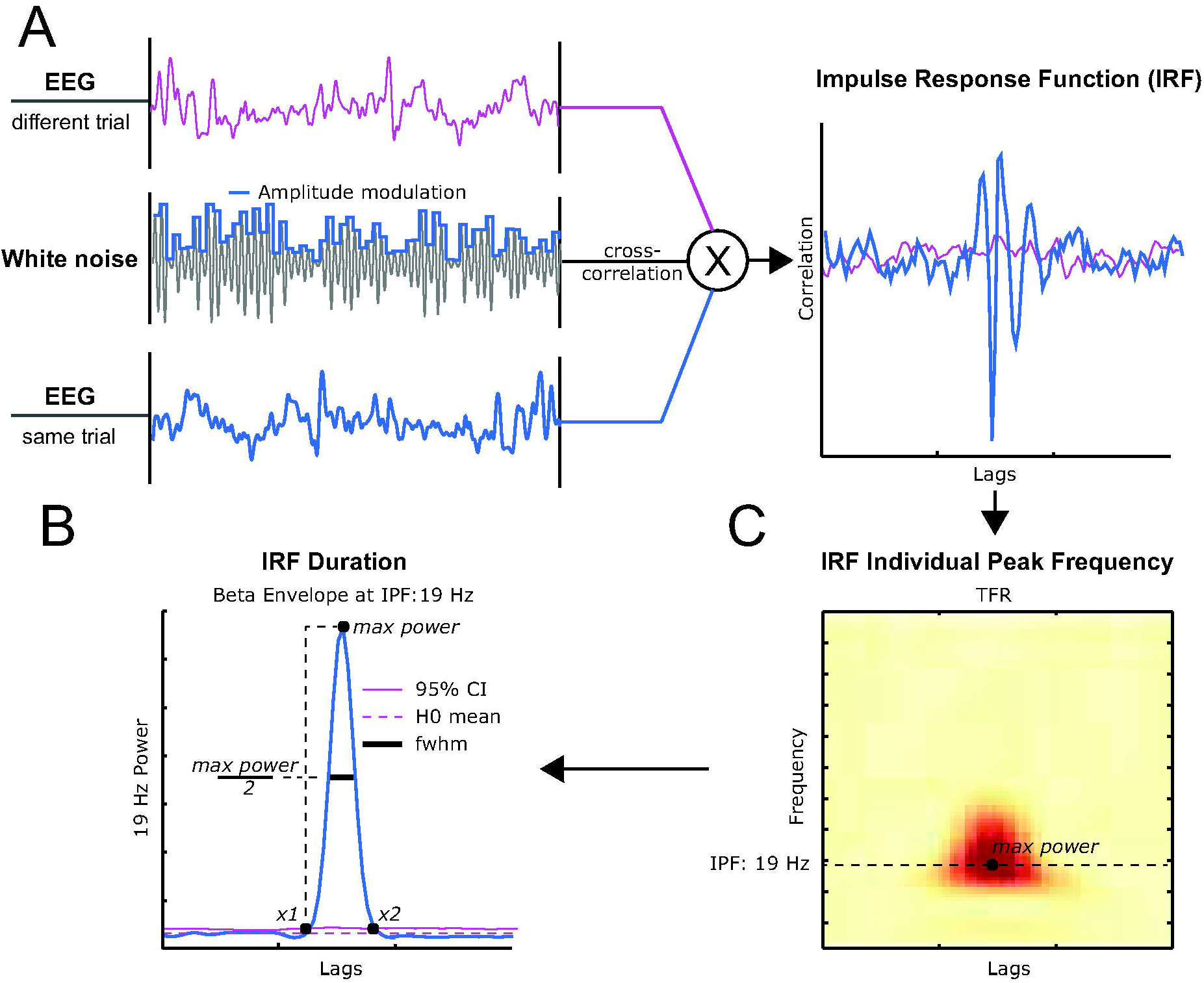
**A**. Cross Correlation Procedure. The cross-correlation between white noise sequences and EEG data from corresponding (same) trials results in the Impulse Response function – the brain response to a single impulse (right panel, blue line). Cross-correlation of white noise sequences with randomly shuffled (different) trials (surrogate IRF) serves as a null hypothesis (EEG and white noise are uncorrelated) and produces an almost flat line (right panel, purple line). **C**. IRF duration was estimated by extracting beta-band power envelope from time-frequency representation (TFR) of IRF signal. **B**. IRF duration was defined as FWHM between beta power peak at fmax and intersects x1 and x2 between the envelope and 95% quantile of the surrogate IRF power distribution at the same frequency.

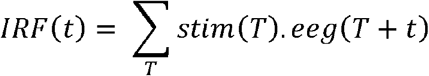

with *stim* and *eeg* denoting z-scored stimulus sequence and corresponding z-scored EEG signal respectively.

#### 4.3 Steady-state Somatosensory Evoked Potentials

Frequency-specific SSSEP responses were isolated using a spatiotemporal source separation method (Cohen, 2022; Cohen & Gulbinaite, 2017), which is based on generalized eigenvalue decomposition (GED) and allows to maximize signal-to-noise ratio of steady-state responses by exploiting information present in inter-channel covariance matrices. Importantly, such approach also allowed us to account for individual differences in somatotopic organization and to separate narrow-band tactile stimulation-related activity from temporally co-occurring broadband movement artifacts in beta band. Thus, instead of analyzing SSSEPs from a subset of electrodes with maximum power at the stimulation frequency, we analyzed a linearly weighted combination of signal from all electrodes.

For each participant and stimulation frequency, a separate spatial filter was constructed by temporally narrow-bandpass filtering (Gaussian filter) the raw data (X) around the stimulation frequency *f* (FWHM = 0.5 Hz) and at the two neighboring frequencies (f ± 1 Hz ; FWHM = 2 Hz). Temporally filtered data (500-6500 ms relative to stimulus onset) was then used to compute covariance matrices: one “signal” matrix (**S** covariance matrix) and two “reference” matrices that were averaged (**R** covariance matrix). The first 500 ms contain evoked potentials that have different sources than SSSEPs (Nangini et al., 2006), and thus were excluded to not compromise the quality of the spatial filter (Cohen & Gulbinaite, 2017). Generalized eigenvalue decomposition (Matlab function eig) performed on “signal” and “reference” covariance matrices returned matrices of eigenvalues and eigenvectors. To increase the robustness of the spatial filters, we applied a 1% shrinkage regularization to the average “reference” covariance matrix. Shrinkage regularization involves adding a percent of the average eigenvalues onto the diagonals of the average “reference” covariance matrix (Cohen, 2022). This reduces the influence of noise on the resulting eigen decomposition. The eigenvectors (column vectors with values representing electrode weights, w) were used to obtain component time series (eigenvector multiplied by the original unfiltered single-trial time series, w^T^ X). The component with the highest signal-to-noise ratio in the power spectra at the stimulation frequency was selected for further analysis (out of total 324 components = 18 subjects x 18 stimulation frequencies, in 264 of cases the 1^st^ component had the highest SNR, in 48 cases the 2^nd^, in 11 cases the 3^rd^, and only 1 case the 4^th^ component). Topographical representation of each component was obtained by left-multiplying the eigenvector by the signal covariance matrix (w^T^S). The obtained topographical maps were normalized and the sign of eigenvector was flipped for 5 subjects that showed spatial peaks opposite to that of the group average. The sign of the components affects only the representation of the topographical maps and has no effect on component time series (Cohen, 2022).

Power at each stimulation frequency was computed using FFT on single-trial component time series in the 500-6500 ms time window (relative to the stimulus onset) and zero-padded to obtain power exactly at the stimulus frequency with 0.1 Hz resolution. The absolute value of FFT coefficients was squared and averaged across trials. To facilitate comparison across SSSEPs elicited by different stimulation frequencies, SSSEP power values were expressed in SNR units:

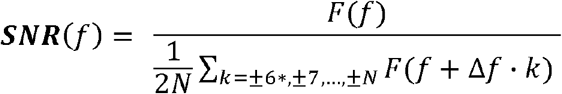

where *N* **=** ± 1.5 Hz (15 bins), excluding 0.5 Hz (5 bins) around the frequency of interest.

For each participant, we plotted SSSEP power and eigenvalues as a function of frequency (frequency tuning curves) and selected the frequency with the highest power as resonance frequency for that participant. In cases when frequency tuning curves contained double peaks, the frequency with the highest eigenvalue was selected. The eigenvalue indicates how well the associated eigenvector separates the “signal” and “reference” matrices and thus can be used as additional selection criterion when comparing components. The statistical significance of SSSEPs was evaluated by repeating the above procedure (constructing spatial filters, obtaining component time series, computing power spectra) using trials in which neither the stimulation frequency nor its harmonics were present. Eigenvector associated with the highest eigenvalue was selected to construct spatial filters expected when no SSSEPs are present. This procedure allowed to determine the SNR values that can be expected when no stimulation at a particular frequency was delivered and to test whether the stimulation protocol induced any significant SSSEP response.

#### 4.4 IRF and ERP Time Frequency Analysis

Time-frequency decomposition for IRFs and evoked potentials in response to single-pulse stimulation were performed via continuous wavelet transformation. The total power of the ERP was calculated by multiplying the power spectrum of single-trial EEG data with the power spectrum of complex Morlet wavelets 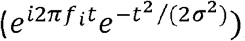, where *t* is time, *f*_*i*_ is frequency that ranged from 2 to 50 Hz in 1 Hz steps, and σ is the width of each frequency band defined as n/(2πf_i_), where n is a number of wavelet cycles). Following publications by Sherman et al. (2017) and Shin et al. (2017) we used 7-cycle Morlet wavelets for all frequencies. Similarly, the spectral content of the IRFs was analyzed by time-frequency decomposition of trial-average IRFs. Power-spectra for IRFs and ERPs (Figure 3 C,F,I) were computed by averaging time-frequency representations in the 0 – 200 ms time window at the subject-specific channel of interest.

**Figure 3.**
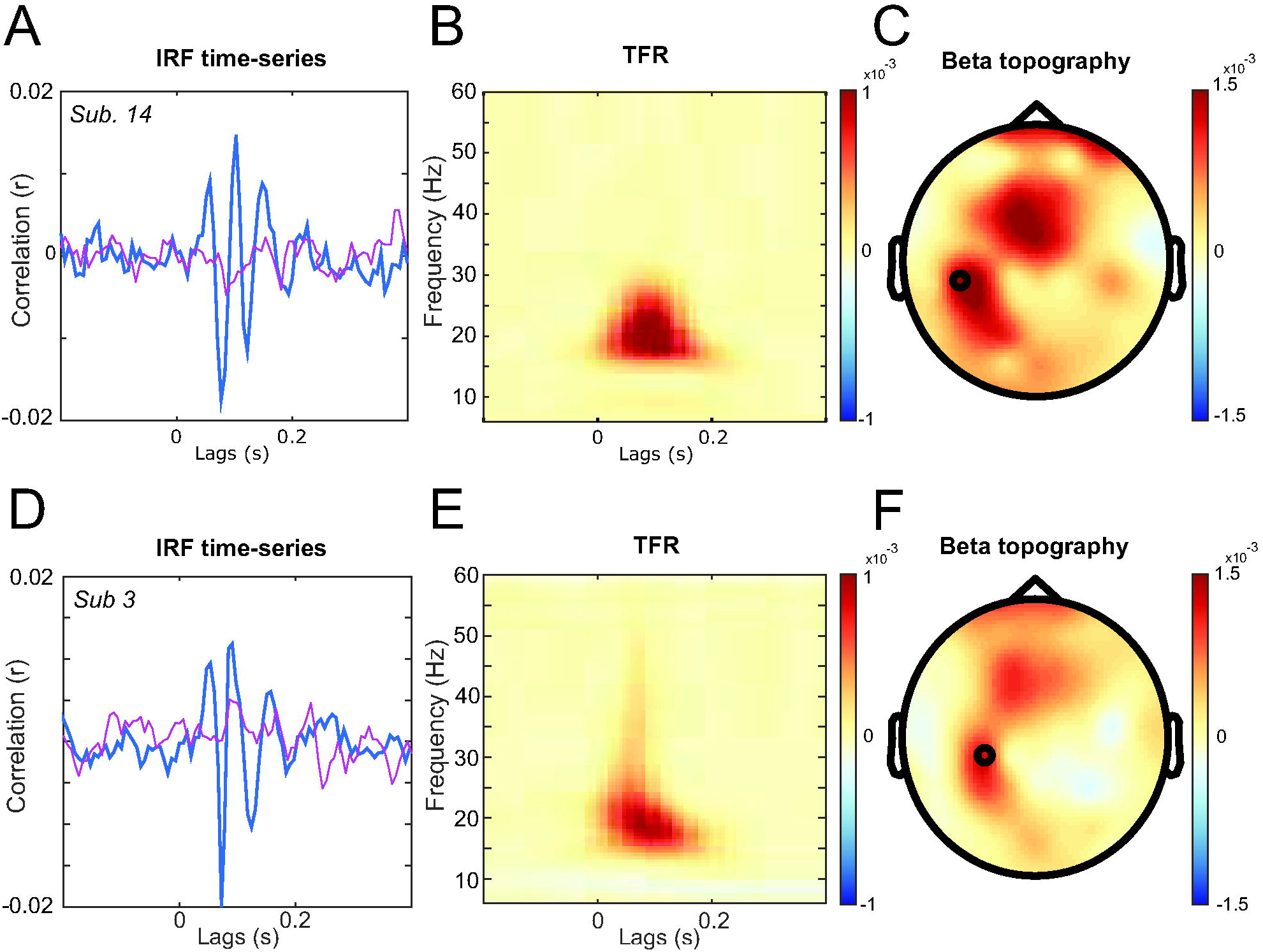
Impulse response functions and their corresponding time-frequency representations from two representative subjects. **A**. The observed IRFs (blue lines) were calculated by cross-correlating tactile WN sequences with the recorded EEG signal. The red lines represent cross-correlation result after randomly shuffling trials (null hypothesis: EEG and white noise are uncorrelated). **B**. Time-frequency representation of the single subject IRF at the individual channel of interest (CP3). **C**. Topography of average IRF beta power (15 Hz to 40 Hz, 0 ms to 200 ms). **D**,**E**,**F**. Same as A,B,C but for a different subject (Channel CP5).

To extract the non-phase-locked part of the EEG response (Figure 4D), we first averaged the time-domain signal and subtracted it from all trials prior to time-frequency decomposition. The phase-locked part of the signal (Figure 4G) was calculated by subtracting non-phase locked power from the total power. Decibel baseline correction was applied to ERP time-frequency representations according to the following formula 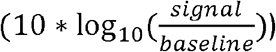 using a baseline window from -400 ms to 100 ms before stimulus onset. Absolute baseline correction was applied to the time-frequency representation (TFR) of the IRF by subtracting the average power values calculated from lags between -400 ms and -100 ms from all frequency bands respectively.

**Figure 4.**
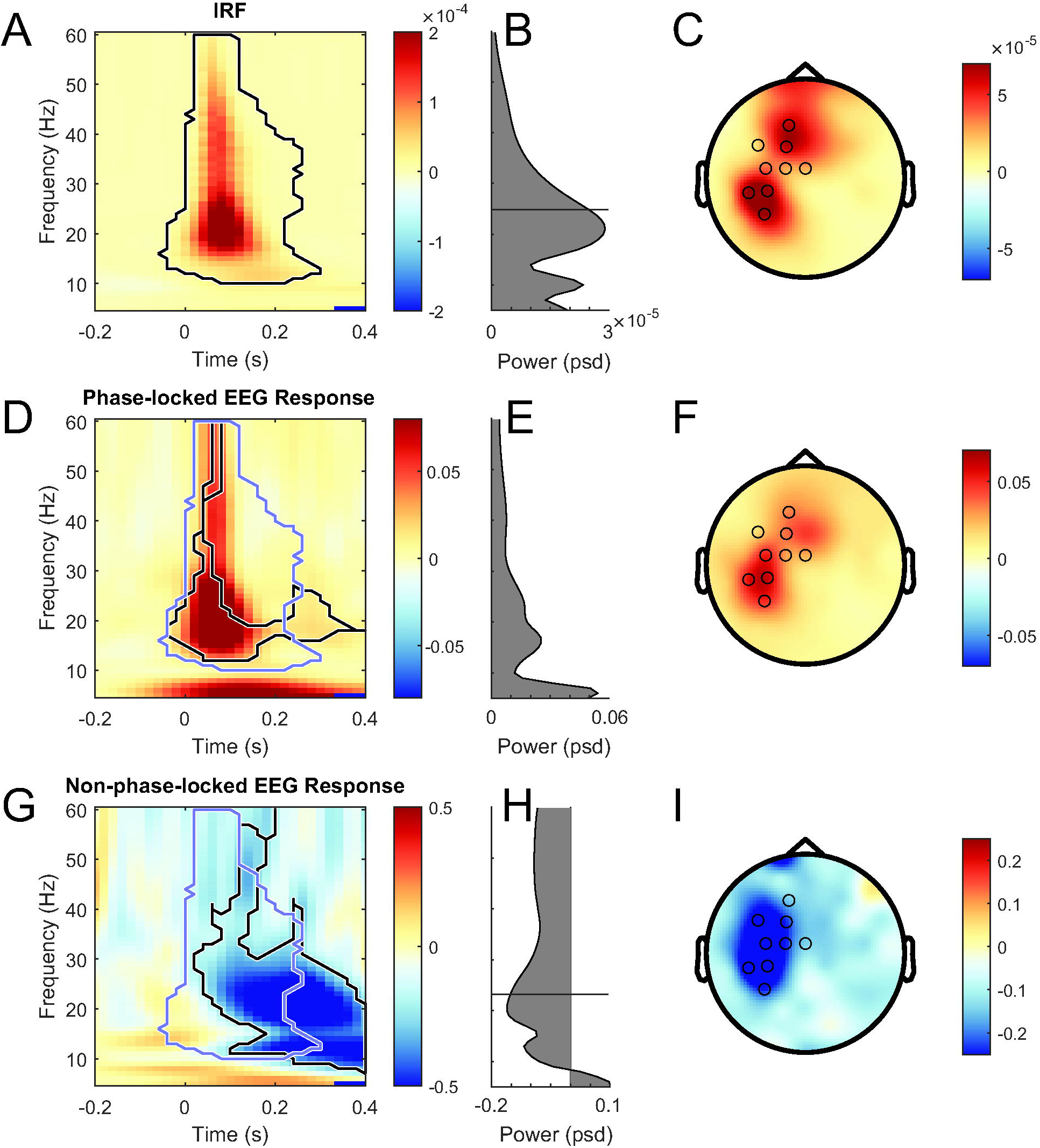
**A**. Grand average time-frequency representation of IRF (averaged over individual channels of interest). Black contours indicate contiguous time-frequency points at which the observed IRF was significantly different from surrogate IRFs at p < 0.05 (corrected for multiple comparisons across time-frequency points using false discovery rate). **B**. Grand**-**average power spectrum of the IRF calculated in the 0 ms to 200 ms time window. Individual subjects showed spectral peaks distributed around a mean of 25.1 Hz. **C**. Topography of IRF beta-band power (15 to 40 Hz, 0 to 200 ms). Black dots indicate the distribution of electrodes that were identified as a subject-specific channel of interest. **D**. Time frequency representation (averaged over individual channels of interest) and beta topography of the phase-locked EEG response (ERP) and **E**. Grand-average power-spectra calculated in the 0-400 ms time window over individual channels of interest for the phase-locked EEG response. **G**. Grand average time-frequency representation of non-phase-locked EEG response. Black contours indicate regions in which contiguous time-frequency points were significantly different from the pre-stimulus baseline at p⍰<⍰0.05 (corrected for multiple comparisons using false discovery rate). Purple contours depict a region, in which the real IRF was significant (the same as black contour in A). **H**. Same as in E. but for non-phase-locked EEG response. Individual subjects showed local minima at frequencies distributed around a mean of 23.1 Hz. **F**,**I**. Topography of beta-band power (15 to 40 Hz, 0 to 400 ms) for the phase-locked and non-phase-locked part of the EEG response.

Given some variability for maximum amplitude IRF channel across subjects (individual differences in anatomy and thus projection on the scalp), subject-specific channels of interest were defined as follows: The time-frequency representation of IRFs was averaged over all channels, and the time-frequency window of interest (0 ms to 200 ms, 15 Hz to 40 Hz) showing increased beta band power was defined based on visual inspection (see Figure 3A and 3G). The channel with the largest power value within this window was selected as the channel of interest for this subject. The frequency at which the largest power was observed was taken as the individual’s IRF peak frequency.

Duration of IRF at the channel of interest was quantified in the following steps. First, we extracted the power envelope of IRF at individuals’ IRF peak frequency (*fmax*) determined from the time frequency representation of IRF (Figure 2C). Then, the IRF duration was defined as the full width at half maximum (FWHM) between the maximum IRF power and intercepts *x1* and *x2*, which were defined as the timepoints where IRF power envelope exceeded the 95% quantile of IRF surrogate power values (null hypothesis distribution).

Time-frequency representations of IRFs were statistically assessed by comparing each individual observed time-frequency power value to the corresponding distribution of power values from the surrogate distribution (Figure 4A). If the observed power exceeded 95% quantile of the surrogate distribution it was considered significant. Time-frequency representations of phase locked and non-phase locked EEG responses were assessed by comparing all participants time-frequency power values (N = 18) to the baseline power values at the corresponding frequency using t-tests. Correction for multiple comparisons was done using false discovery rate. Before comparison, baseline power values were averaged between -400 ms and -100 ms for each individual frequency. We only considered the largest cluster of significant time-frequency points for further analysis.

#### 4.5 Spontaneous beta burst analysis

Spontaneous beta bursts were extracted from the -500 ms to 0 ms baseline activity from all 3 paradigms (white noise stimulation, SSSEP, and single-pulse stimulation). Single-trial time frequency representations were calculated at subject-specific IRF channels of interest as described in section 3.4. Power thresholds for beta burst detection were set to be 3 times the standard deviation of all beta-band (15 Hz to 40 Hz) power values calculated over all trials. Before calculating the standard deviation, we removed 1% of the largest power values to reduce noise stemming from high-power artifacts. We used Matlab’s *imregionalmax* function to detect peaks (between 15 Hz and 40 Hz) in the time-frequency representations that exceeded the power threshold. From the bursts that matched our selection criteria we calculated individual frequency histograms. Individual peak frequencies were defined as the highest peak in the frequency histograms.

## Results

The power of ongoing (non-phase-locked) beta oscillations is generally decreased in response to tactile stimulation and motor actions (Hari et al., 1997; Neuper & Pfurtscheller, 2001; Parkkonen et al., 2015; Salenius et al., 1997; Salmelin & Hari, 1994). More recent work capitalizing on single-trial analysis has demonstrated that spontaneous beta bursts occurring in close temporal proximity to tactile targets deteriorate stimulus detection (Little et al., 2019; Lundqvist et al., 2018; Shin et al., 2017). These and other findings have led to the hypothesis that somatosensory beta oscillations serve as an inhibitory brain rhythm that is generally absent during periods of increased processing demands for incoming information (Karvat et al., 2021). Given the aforementioned reports on burst-like nature of beta oscillations and previous findings of the impulse response function in the visual domain (VanRullen & Macdonald, 2012), we set out to test if the somatosensory system also responds to white noise (WN) vibrotactile stimulation with a rhythmic impulse response function (IRF) – a transient increase in beta-band power. We first characterized the frequency and duration of the IRF in response to WN, and then we tested whether the properties of the IRF are related to: 1) the resonance properties of the somatosensory system measured in a SSSEP paradigm; 2) dynamic changes in beta power in response to single-pulse stimulation; and 3) spontaneous beta bursts occurring during the intertrial interval (a proxy for the resting state beta-band activity).

### 1. Random Noise Stimulation reveals oscillatory Impulse Response Function

Cross-correlation of WN tactile stimulation sequences with concomitantly recorded EEG signal revealed an oscillatory response present for lags up to ∼200 ms – the impulse response function (IRF) of the somatosensory system (Fig. 3). Topographically, the IRF amplitude was maximal over the left parietal and frontal channels. The time-frequency representation of IRF showed that these fluctuations in response to tactile impulses were constrained to the beta band. Spatial and frequency characteristics of the somatosensory IRF were similar to previous reports on vibrotactile stimulation, with the highest beta power observed contralaterally to the stimulated hand (Bardouille et al., 2010; Bardouille & Ross, 2008; Colon et al., 2012; Langdon et al., 2011; Nangini et al., 2006; Spitzer et al., 2010; Vlaar et al., 2015), thus indicating a genuine somatosensory response (Figure 4B).

To test whether the beta oscillatory response to WN tactile stimulation was merely a result of a general increase in beta power in response to tactile stimulation, we repeated the same analysis steps, however this time we cross-correlated WN sequences with random EEG trials 10.000 times and re-performed spectral analyses. We statistically compared the observed average power values at every time-frequency point to the distribution of power values from the surrogate distribution. Real IRF power values were significantly higher for lags between 0 and ∼200 ms and frequencies between 10 and 60 Hz. The power spectra calculated by averaging over all timepoints between 0 ms and 200 ms showed a distinct peak at 20.1 Hz which was not present in the surrogate distribution (Figure 4C).Taken together, this initial assessment suggests that the somatosensory system responds to tactile stimulation with an oscillatory beta burst that is: 1) phase-locked to tactile stimulation, and 2) specific to the exact white noise sequence that was presented on that particular trial.

Next, we quantified individual participants’ IRF peak frequency to estimate the length of individual IRFs and to compare its spectral properties to other beta signatures. Analyzing the maxima in participants TFRs at individual channels of interest allowed us to identify individual peak frequencies ranging from 18 to 39 Hz (mean = 25.1, std = 6.12). These frequencies are well within the beta band although IRF peak frequency of two subjects (39 Hz) neared the upper edge of classically defined beta band.

### 2. Duration of IRFs

Visual IRFs can be observed for up to 1 second, and last on average 4 to 5 alpha cycles (Brüers, 2017). We were curious if the short (∼200 ms) duration of tactile IRFs as compared to the visual ones was a result of oscillatory frequency difference between tactile and visual IRFs: Albeit temporally shorter, tactile IRFs might oscillate for the same number of cycles as their visual counterpart. We therefore quantified the length of each subject’s tactile IRFs (at the channel of interest) based on the power envelope at individual peak frequency. The average duration (full width half maximum relative to 95% quantile surrogate distribution) of individual tactile IRFs was 116.58 ms (SD = 19.13). We used the following formula to calculate the number of cycles for a given individual peak frequency 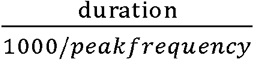. Average length in cycles (calculated from FWHM) across subjects was 2.81 (SD = 0.3544). Thus, while tactile IRFs are slightly shorter than visual IRFs, they closely match the length (3 cycles) of spontaneous beta bursts reported in previous work (Little et al., 2019; Lundqvist et al., 2016).

### 3. Non-phase-locked beta power and IRF share spectral properties

To directly compare the spectral properties of IRFs with beta-band activity in response to traditional sparse stimulation protocols, we performed time-frequency analyses of ERP data. We separately analyzed non-phase-locked and phase-locked activity (see Methods section for details). Similar to previous reports (Hari et al., 1997; Neuper & Pfurtscheller, 2001; Parkkonen et al., 2015; Salenius et al., 1997; Salmelin & Hari, 1994), non-phase-locked beta power decreased significantly as a result of tactile stimulation (Figure 4G). This decrease was most pronounced over the left parietal/frontal channels (Figure 4I). For each participant, we identified the peak frequency at which the strongest decrease in beta power occurred by detecting power troughs following the stimulus onset (Figure 4H). Individual minima occurred at frequencies ranging from 19 to 33 Hz (mean = 23.1 Hz, std = 3.34 Hz). To test if the non-phase-locked beta signatures and IRFs might share a common neural generator, we correlated individual beta trough frequencies. Spearman correlation between IRF and decreases in non-phase-locked beta power revealed a significant relationship (r = 0.55, p = 0.0216, Figure 7) indicating that both beta signatures might rely on a common oscillatory generator.

### 4. Phase locked beta power and IRF share spectrotemporal properties

The phase-locked part of the EEG response showed a broad band increase in power over somatosensory areas (Figure 4D and 4F). Statistical analysis of the time-frequency representations revealed a cluster of significantly increased power across a large frequency band (15 Hz to 60 Hz) between ∼0 ms and ∼200 ms that was followed by a more frequency specific increase in the 15 to 30 Hz range (∼200 ms to 400 ms; Figure 4D).

Intriguingly, the TFR’s of phased locked ERPs closely resembled the TFRs of the IRFs (Figure 4A,D). Related, we found beta-band topographies to be highly similar (Figure 4C,F). To test if IRFs constitute the initial part of the commonly investigated ERPs we sought to identify spectral peaks in the TFR of phase locked ERPs and compare them to the IRF peak frequency. We could identify spectral peaks in all subjects (mean = 16.28 Hz, std = 5.04), however for 4 of these subjects the peaks did not fall within the significant cluster (Figure 4D) but were significantly lower (6 to 12 Hz). Furthermore, the peak frequencies did not correlate with IRF peak frequencies (r = -0.21, p = 0.4, Spearman). This null finding reflects the fact that time-frequency representation of ERP is dominated by the low frequencies (Figure 4D), and frequency-based analyses are suboptimal to uncover short-lived high-frequency spectral components. We thus performed time-frequency decomposition of ERPs and directly compared time-frequency representations (TFR) of IRFs and ERPs. Our rationale was as follows: If phase locked ERPs contain IRFs and if IRFs vary between participants then within-participant ERP and IRF correlation should be higher as compared to between-participant ERP and IRF correlation.

All time-frequency values of an individual IRF cluster (Figure 4A, Figure 5A) were correlated (Pearson r) with the corresponding time-frequency values of the ERP (Figure 4D, Figure 5A) within and between subjects yielding a correlation matrix, in which diagonal as compared to off-diagonal correlation values should be significantly higher (Figure 5B). We averaged all row and column vectors corresponding to individual diagonal entries and compared the resulting values to the diagonal using a t-test (Figure 5CD). Time-frequency representations of individuals IRF were significantly higher correlated with the same individuals ERP as compared to the ERP of all other individuals (Figure 5D, t(17) = 2.28,p = 0.035, effect size d = 0.437). This indicates that some spectral components of the phase locked ERP might in fact be oscillatory IRFs, embedded in the ERPs low frequency components

**Figure 5.**
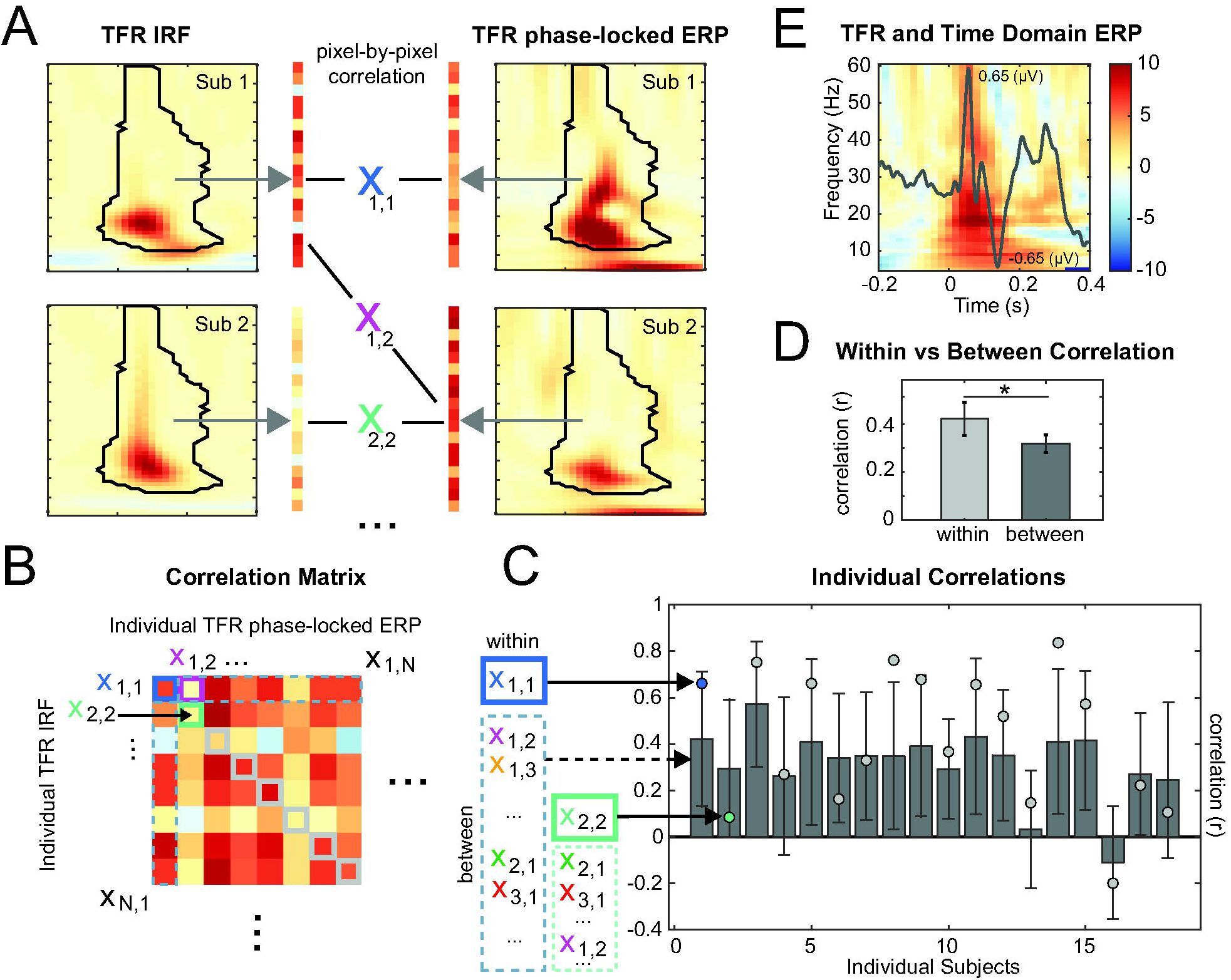
**A**. Within and between subject correlation of IRF and phase-locked EEG Response (ERP) time frequency representations. Pixel values within the black outlined area were correlated using Pearson correlation. **B**. Resulting Correlation Matrix. **C**. Individual within-subject correlation values (light grey circles) and between subject correlations (dark grey bars, error bars indicate 95% confidence intervals). **D**. T-test between within-subject and average between subject correlation values. E. Characterization or ERP in Time and Frequency Domain (Group Average, Channel CP5).

### 5. Somatosensory resonance frequency matches the frequency of somatosensory IRFs

From the linear dynamical systems perspective, the frequency response of a system – the system’s response to a sinusoidal input with variable frequency – should be similar to the frequency transform of its impulse response function. To test this hypothesis and determine subject-specific resonance frequencies of the somatosensory cortex we employed sine-wave stimulation in 12-39 Hz frequency range (SSSEP paradigm). Steady-state evoked potentials were extracted using a spatiotemporal source separation method that allows to linearly combine information from all the channels instead of analyzing SSSEPs from channels with maximum power at the stimulation frequency (Cohen & Gulbinaite, 2017).

As expected, vibrotactile right index finger stimulation elicited maximal response over the contralateral somatosensory cortex (Figure 6A). Although for all participants SSSEPs were strongest over the frontocentral left-hemifield channels, substantial across-subject variability was evident. To accommodate for individual differences in SSSEP scalp projections, which are likely caused by anatomical differences, we linearly combined information from all the channels by computing spatial filters (channel weights) that optimally isolate frequency-specific SSSEPs (see Methods section for details). This was done separately for each participant and each stimulation frequency.

**Figure 6.**
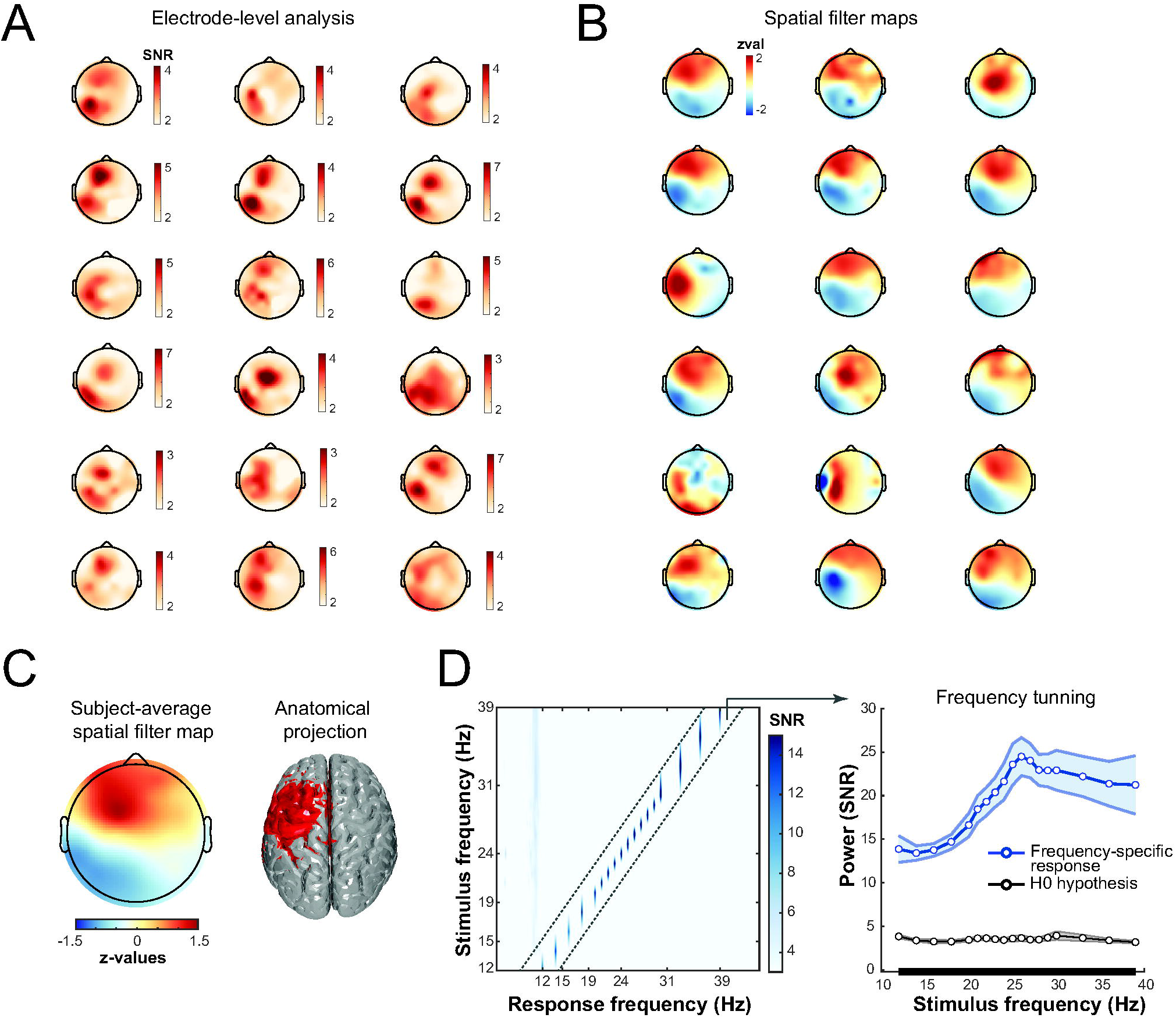
Steady-state somatosensory potential analysis. **A**. Electrode-level analysis results. Each topographical map represents single-subject power (expressed in SNR units) averaged across all stimulation frequencies (12-39 Hz, 18 frequencies). Note the individual variability in scalp projections of the SSSEP responses. **B**. Topographical representation of single-subject spatial filter corresponding to each subject’s resonance SSSEP frequency. Note the high similarity across subjects and dipolar appearance of the components. **C**. Subject-average normalized spatial filter weights depicted in B (left) and the associated putative anatomical generators of SSSEPs. **D**. Subject-average SSSEP power expressed in SNR units plotted as a function of vibrotactile stimulation frequency (left panel): Input frequencies are depicted on the y-axis and output frequencies on the x-axis. The brighter colors indicate stronger response, with maximal responses observed, as expected, at the stimulus frequency. Right panel depicts SSSEP power at stimulation frequencies (diagonal from the matrix of power spectra on the left), with the resonance peak at 26 Hz. Black line “H0 hypothesis” represents SSSEP amplitude at each of the tested frequencies on trials when stimulation at that frequency was not delivered and can be expected as a result of overfitting. Black line on the x-axis indicates that SSSEPs at all frequencies were statistically significant at p< 0.001 (corrected for multiple comparisons across frequencies using cluster-based permutation testing).

Figure 6B depicts normalized spatial filter weights for the resonance stimulation frequencies averaged across participants. The putative anatomical estimate of SSSEPs determined from the subject-average spatial filter was localized, as expected, over the left somatosensory cortex (Figure 6B).

The subject-average response power (expressed in SNR units) plotted as a function of stimulus frequency showed the strongest responses at the stimulation frequency (diagonal). Harmonic responses that are sometimes reported for rhythmic vibrotactile stimulation paradigms are not visible here, because spatial filters were designed to isolate responses to narrow-band rhythmic stimulus and suppress temporally co-occurring activity at other frequencies. All stimulation frequencies elicited statistically significant SSSEPs (for all stimulation frequencies p < .001, corrected for multiple comparisons across frequencies using cluster-based permutation testing), as indicated by the small SNR at each of the 18 tested frequencies on trials when stimulation frequency was not present (Figure 6D; see Methods section for details). The highest amplitude SSSEPs, or resonance frequency, on average was around 26 Hz (22-39 Hz range, SD = 5.32 Hz). These findings are consistent with previous reports that used a “best” electrode(s) approach and reported a resonance of human somatosensory cortex in the 20-26 Hz range (Müller et al., 2001; Snyder, 1992; Tobimatsu et al., 1999).

We extracted individual peak resonance frequencies from all subjects and compared them to the frequency of IRFs and non-phase locked beta power decreases following stimulation. Resonant frequencies were correlated significantly with peak frequencies of IRF and stimulus related beta power decreases (p = 0.0003 and 0.0195 respectively, Spearman correlation, Figure 7). This result suggests that the frequency of increasing and decreasing beta power signatures during stimulus processing might be determined by the fundamental resonance frequency of the somatosensory system.

**Figure 7.**
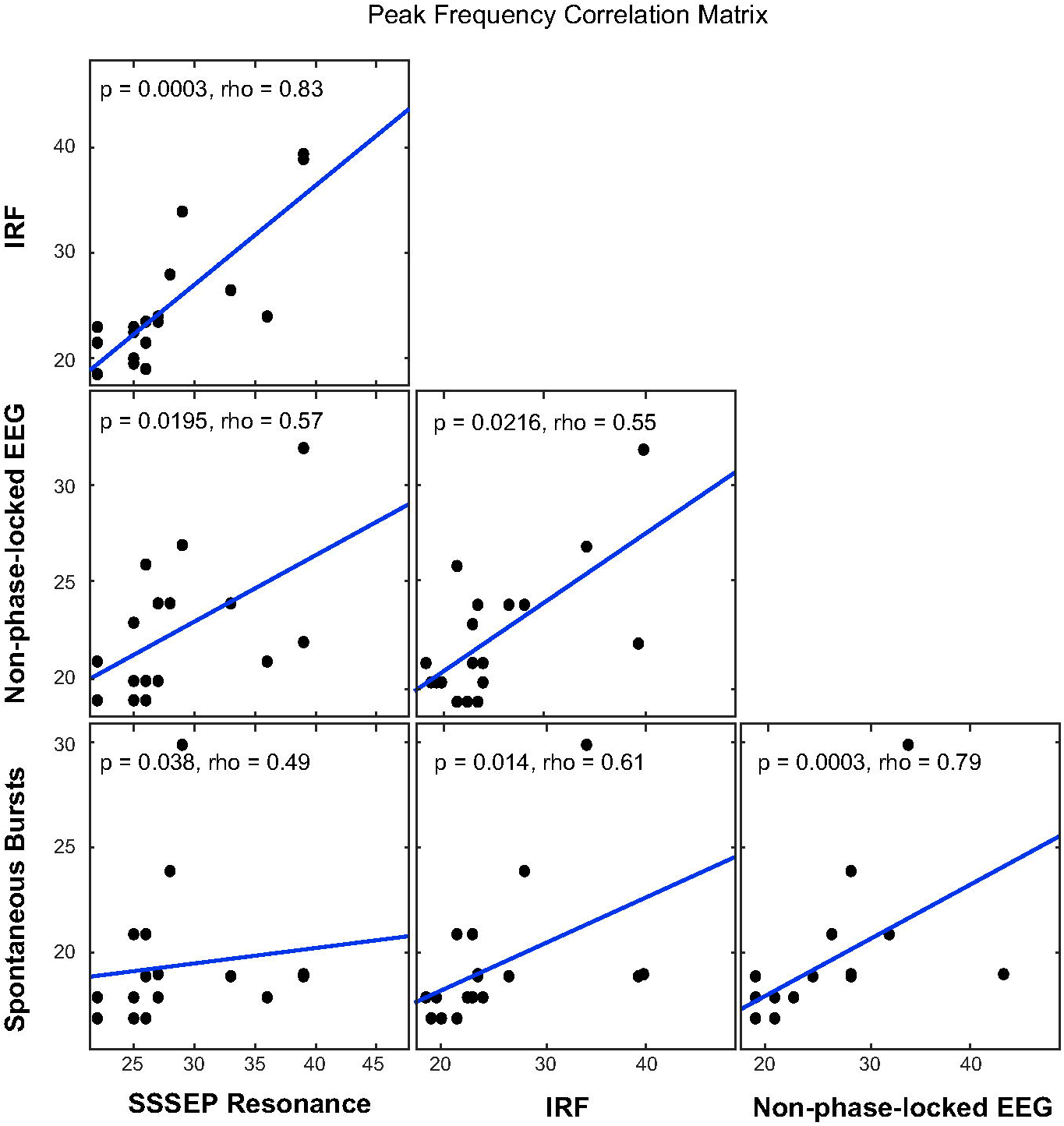
Peak Frequency Correlation analysis. Spearman correlations were calculated between individual peak frequencies of IRFs, non-phase-locked EEG response and spontaneous Bursts measured during baseline. Correction for multiple comparisons was performed using FDR.

### 6. Spontaneous beta bursts and IRFs share spectral properties

Recent studies focused on beta-band oscillations at a single-trial level and revealed that instead of sustained beta oscillations present in trial-average time-frequency representations, beta oscillations are transient burst-like events (Little et al., 2019; Lundqvist et al., 2018; Shin et al., 2017). The duration (3 cycles), frequency range (15 – 30 Hz) and location (somatosensory cortex) of the reported transients is highly similar to the spectro-temporal properties of IRFs that we report here (mean = 2.8, frequency range 18 – 39 Hz). To investigate if the two share neural generators, we quantified average beta burst frequency in our dataset and compared them to individual IRF peak frequencies. Beta burst analysis was performed in the pre-stimulus window (−500 - 0 ms) from all trials (SSSEP, white noise and single-pulse ERP trials). Bursts were detected from single trial TFR’s by using a threshold of 3 standard deviations (see Methods section for details). We could identify peaks in all subjects in the 17 to 30 Hz range (mean = 19.44, std = 3.06). Individual burst peak frequencies were significantly correlated with peak frequencies of SSSEP resonance (p = 0.038), IRF (p = 0.014) and non-phase locked beta power decreases (p = 0.0003) (Spearman correlation, Figure 7). Our analysis provides evidence that several well investigated beta phenomena are all strongly correlated, and potentially reflect activity of the same neural generators, linked to the fundamental resonant properties of the somatosensory system.

## Discussion

The visual system’s response to an impulse of light is oscillatory (“perceptual echo”) and strongly correlates with the endogenous alpha rhythm (VanRullen & Macdonald, 2012) – a feature revealed by cross-correlating stimulus white noise sequences with the corresponding EEG trials. These findings suggested that response to visual stimulation is phase-locked to the stimulus and may play an active role in processing stimulus specific information. Here we extend these findings to the somatosensory domain by showing that the somatosensory cortex responds to tactile stimulation with a phase-locked 3-cycle long reverberation in the beta band akin to a burst. These stimulus-locked beta bursts were significantly correlated in peak frequency with the non-phase-locked decrease in beta power following tactile stimulation, the individual participants’ tactile resonance frequency (determined in SSSEP paradigm) and the frequency at which spontaneous beta bursts occur during rest. Our findings suggest a common oscillatory generator underlying all beta signatures characterized in this study, and reveal a supramodal mechanism in sensory processing.

### 1. Phase-locked beta bursts in response to tactile stimulation

The cross-correlation between tactile white noise (WN) sequences and the corresponding EEG signal revealed oscillatory signatures that showed strong power in the beta band (∼22 Hz) over somatosensory cortex. Theoretically, the IRF revealed using cross-correlation should resemble a classical time-domain averaged ERP (Polge & Mitchell, 1970). Our findings, however, revealed significant differences. The spectral content of the phase locked part of the ERP was dominated by frequencies in the 1-15 Hz range (Figure 4D). The IRF, on the other hand, showed very little low-frequency components and contained a burst-like increase in beta power, that lasted for ∼3 cycles.

The most surprising finding, in the context of previous research, is the phase-locked nature of somatosensory echoes. In contrast, previous reports have highlighted spontaneous occurrence of beta-band bursts in the empty intervals before target detection (Shin et al., 2017), working memory delay intervals (Lundqvist et al., 2018) or movement preparation and response errors (Little et al., 2019). This precise time-locking strongly suggests that somatosensory echoes are directly involved in the processing of tactile stimuli and do not likely reflect secondary processes related to movement planning or working memory maintenance. Which functional role could such an oscillatory signature play? Communication through coherence assumes precise phase alignment to be the key factor in information transfer (Bastos et al., 2015; Fries, 2015). While high frequency oscillations, such as gamma, usually increase in power following stimulation, low frequency rhythms, such as alpha or beta, decrease in power in these critical moments of information transfer (Figure 3D). Our finding of phase-aligned low frequency beta oscillations during this critical early stage of processing indicates that beta might be fundamentally involved in bottom-up transfer of tactile information.

### 2. The relationship between ERPs and IRFs

Tactile stimulation of the skin or electrical stimulation of the median nerve is generally accompanied by two EEG signatures in the somatosensory-system: One is a broad-band, phase-locked increase in power, and the other is a non-phase locked initial suppression of mu and beta rhythm followed by an increase (so called “beta rebound”, Hari et al., 1997; Neuper & Pfurtscheller, 2001; Parkkonen et al., 2015). It has been a matter of debate if the initial phase-locked broad-band EEG response (ERP) is generated, at least partially, by phase reset of ongoing oscillations (Sauseng et al., 2007). Somatosensory evoked potentials (SEPs) commonly show two early components (N20 and N60), the interpeak difference between which is ∼40 ms and variable across individuals (Niedermeyer & da Silva, 2005). The interpeak timing could potentially correspond to a 25 Hz oscillation and is similar to average somatosensory IRF frequency, as well as the peak of somatosensory frequency tuning curves reported here. High correspondence in spectro-temporal characteristics between these different electrophysiological measures of somatosensory system (SEPs, somatosensory IRFs, peak of somatosensory tunning curves) indicates that they all capture resonance characteristics of the underlying neural network.

While ERPs could also be considered IRFs (Zaehle et al., 2010), there are some important differences between the two: First, ERPs are responses to sparse tactile stimuli whereas IRFs reflect responses to continuous increments and decrements of tactile stimulation intensity. Second, the white-noise stimulation technique is much more efficient: While in the computation of ERPs, one sample per ERP-timepoint is collected per trial, the cross-correlation considers all samples to calculate similarity between the signals. Third, short and sparse presentation of stimuli used in experiments strongly differs from natural tactile stimulation and perception. Natural perception is much more continuous and incoming information fluctuates dynamically at multiple frequencies. Thus, IRFs might be a more accurate characterization of neural EEG correlates of sensory processing. Notably, calculating the IRF via cross-correlation assumes a linear relationship between input and output which is likely violated in neural processing, e.g. due to the involvement of multiple mechanoreceptors that bandpass filter the input frequencies (Handler & Ginty, 2021). The IRF will hence only capture a linear part of brain response and ERP and IRF might therefore reflect different aspects of the same underlying physiological process. Furthermore, although arguably non-linear, the visual IRF does, to a certain degree, capture functionally relevant components that can predict perceptual outcomes (Brüers & VanRullen, 2018).

### 3. Characterizing the resonant properties of the somatosensory system

Experiments using rhythmic stimulation have shown that the somatosensory system responds more strongly to beta-band frequency stimulation (Müller et al., 2001; Snyder, 1992; Tobimatsu et al., 1999). Using an SSSEP paradigm in combination with a spatiotemporal source separation method, we investigated the resonance properties of the somatosensory system and compared it to the spectral characteristics of tactile IRFs, and spontaneous beta bursts occurring during inter-trial intervals. In line with previous reports, we found the mean resonance frequency (highest SSSEP power) to be around 25 Hz ((Snyder, 1992): 26 Hz, (Müller et al., 2001): 27 Hz, (Tobimatsu et al., 1999): 21 Hz). Statistical analysis revealed a significant positive correlation between individual SSSEP peak frequencies, individual IRF peak frequencies and spontaneous beta burst frequencies. We interpret these consistent correlations as evidence for a common neural generator that is primarily determined by the resonance properties of the somatosensory system.

A remaining question is to which extend peripheral mechanoreceptors could give rise to the observed beta oscillations. The time-varying stimuli we used (150 Hz carrier frequency and 12-39 Hz amplitude modulated sine-wave stimulation) likely activate two types of mechanoreceptors responsive to vibrations of the skin: Pacinian corpuscles are most sensitive to high frequency 100-400 Hz vibrations and Meissner’s corpuscles, which respond optimally to 40-60 Hz vibrations (Handler & Ginty, 2021). The here reported rhythmic responses (IRF: 25 Hz, Resonance: 26 Hz) fall below the optimal (resonance) frequencies of these. The amplitude modulation contained frequencies between 0 Hz and 75 Hz and could therefore potentially drive Meissner’s corpuscles. These receptors however have phasic responses and only fire upon initial contact with an object to the vibrations caused by the movement between object and skin (Abraira & Ginty, 2013). As we discarded the initial 500 ms and the last 1000 ms of each individual epoch we find it unlikely that the responses of these receptors could have given rise to the specific beta band oscillations observed in the EEG. Last our stimulation could have activated Merkel’s discs as they respond to sustained point pressure. The firing rate of these cells lies in the 60 Hz range and is hence also unlikely to lead to consistent 20 Hz oscillatory activity in the Scalp EEG signal (Iggo & Muir, 1969). We conclude that the observed rhythms most likely hallmark active processing and originate from cortical or subcortical regions.

### 4. Impulse response functions as multimodal signatures of cortical processing

Beta oscillations have been associated with a multitude of processes such as motor stiffening (Baker et al., 1997), attention (Buschman & Miller, 2007; Lee et al., 2013), status quo maintenance (Engel & Fries, 2010), sensorimotor integration (Androulidakis et al., 2006, 2007; Baker, 2007; Gilbertson et al., 2005), working memory (Haegens et al., 2017; Lundqvist et al., 2018; Schmidt et al., 2019; Spitzer & Haegens, 2017), time perception (Baumgarten et al., 2015; Wiener et al., 2018), and top-down communication (Bressler & Richter, 2015; Buschman & Miller, 2007; Cannon et al., 2014; Lee et al., 2013). Importantly, these theories propose an active role of beta oscillations in temporally organizing incoming information. The stimulus specific nature and strict temporal alignment of beta-band somatosensory IRFs as we demonstrate here, thus could provide a rhythmic temporal structure for these processes.

Two recent studies have implicated the IRFs in regularity learning and predictive coding processes. Repeated presentation of a single visual WN sequence leads to a gradual increase in oscillatory alpha power in the corresponding IRF (Chang et al., 2017). This increase is persistent even when the repetition is interleaved with a novel WN sequence. The authors hypothesized that, in line with the idea that the visual system dynamically encodes visual sequences, the IRF is a signature of this regularity learning mechanism. The authors discussed the possibility that IRFs might reflect a rhythmic updating of predictions about expected stimulation patterns, thus giving rise to 10 Hz oscillations. This line of arguments was later supported by a simple computational predictive coding model capable of generating physiologically plausible IRFs, which propagated as travelling waves along the visual hierarchy (Alamia & VanRullen, 2019). Their results were verified in human EEG studies that showed similar travelling wave dynamics that altered their direction depending on whether visual input was present or not (Pang et al., 2020). Together these results suggest that visual IRFs might reflect predictive coding processes that continuously generate and update predictions about the incoming information at an alpha rhythm (Clark, 2013; Friston, 2005; Linares et al., 2009; Seth, 2014). Their locus of generation might therefore lie in the interactions between different columns in the visual hierarchy hypothesized to implement predictive coding computations, potentially through canonical microcircuits (Bastos et al., 2012).

The fact that similar oscillatory IRFs are found in two distinct modalities (tactile and visual) might reflect a general process that is fundamental to cortical processing. Previous work has attempted to quantify auditory IRFs and found ERP like responses (Lalor et al., 2009; Lalor & Foxe, 2010). Notably however, oscillatory IRFs have not been found in the auditory domain, potentially due to architectural differences between visual and auditory cortical hierarchies (İlhan & VanRullen, 2012). It was also suggested that the auditory system could rely on fundamentally different strategies to process sensory inputs as auditory information, in contrast to visual and tactile information, is defined in terms of temporal fluctuations across multiple frequency bands (VanRullen et al., 2014). It might well be that auditory IRFs could be revealed using more naturalistic stimuli such as speech, as speech contains multifrequency “perceptual units” (in the words of Giraud & Poeppel, 2012) that range from low-frequencies (syllabic rates; 1-8 Hz) to high-frequencies (phonetic features; 30-40 Hz; Dikker et al., 2020).

## Conclusion

In conclusion, we find that similarly to the visual domain, the impulse response function of the somatosensory system is oscillatory, with a maximum in the beta band. This means that the response to a single tactile impulse is reverberated in the beta band for around 3 cycles. These reverberations significantly correlate in frequency with the resonance frequency of the somatosensory system and spontaneously occurring beta bursts during rest, pointing to a common oscillatory generator. Our findings provide evidence that the beta rhythm is actively involved in tactile stimulus processing. Furthermore, the existence of rhythmic IRFs in the somatosensory domain as well as the visual domain support the idea that they reflect a more basic, modality-agnostic signature of sensory processing.

## Acknowledgments

The authors thank Barkın İlhan for the Matlab code used as a basis to generate stimulus sequences and microcontroller-based device used in İlhan and VanRullen (2012) study, which was connected to vibrotactile electromagnetic solenoid-type actuator and used to deliver stimuli in this study.

